# Residue conservation and solvent accessibility are (almost) all you need for predicting mutational effects in proteins

**DOI:** 10.1101/2025.02.03.636212

**Authors:** Matsvei Tsishyn, Pauline Hermans, Fabrizio Pucci, Marianne Rooman

## Abstract

**Motivation:** Predicting how mutations impact protein biophysical properties remains a significant challenge in computational biology. In recent years, numerous predictors, primarily deep learning models, have been developed to address this problem; however, issues such as their lack of interpretability and limited accuracy persist.

**Results:** We showed that a simple evolutionary score, based on the log-odd ratio (LOR) of wild-type and mutated residue frequencies in evolutionary related proteins, when scaled by the residue’s relative solvent accessibility (RSA), performs on par with or slightly outperforms most of the benchmarked predictors, many of which are considerably more complex. The evaluation is performed on mutations from the ProteinGym deep mutational scanning dataset collection, which measures various properties such as stability, activity or fitness. This raises further questions about what these complex models actually learn and highlights their limitations in addressing prediction of mutational landscape.

**Availability:** The RSALOR model is available as a user-friendly Python package that can be installed from the PyPI repository. The code is freely available at https://github.com/3BioCompBio/RSALOR.

## Introduction

Accurately estimating the fitness of variants is essential both from a biomedical perspective, to deepen our understanding of the mechanisms underlying pathogenesis [1], and from a biotechnological perspective, to improve protein engineering approaches [2]. As a result, an impressive number of computational tools have been developed over the last decade to predict the effects of variants on different protein biophysical properties [3, 4, 5]. These tools are characterized by a wide range of architectures, ranging from simple linear models applied to a few features to complex deep learning methods.

In recent years, the field has witnessed an even more remarkable growth, with the emergence of protein language models (pLMs), which have significantly advanced variant effect prediction [5, 6]. However, these deep learning techniques also come with notable drawbacks. Their immense number of parameters requires extensive training, making them computationally expensive and more prone to overfitting. While there are methods to mitigate overfitting, it remains challenging to disentangle true biophysical properties from unwanted biases and to achieve good generalizability. Additionally, their inherent complexity often makes it nearly impossible to extract meaningful biophysical insights from their predictions.

An alternative approach is to introduce simple prediction models with biological significance that retain the same accuracy as these more complex approaches. Following this direction [7], we recently showed how a simple evolutionary score, when scaled by the relative solvent accessibility (RSA) of the mutated residues, can accurately predict changes in folding free energy upon mutations. The model is extremely simple, interpretable and performant, and has no free parameters to optimize. Its score reflects the impact of a variant on protein fitness, which is broadly defined as the ability of the protein to effectively perform its biological function. As such, it is related not only to stability but also to other protein properties. We thus extended the analysis of this model, called RSALOR, by providing additional evidence of its accuracy across a broader range of protein biophysical properties, including stability, binding affinity, organismal fitness, activity, and expression, using ProteinGym [6], a widely used benchmark dataset specifically designed for this purpose. A graphical representation of the model is provided in Fig. 1.

**Figure 1.**
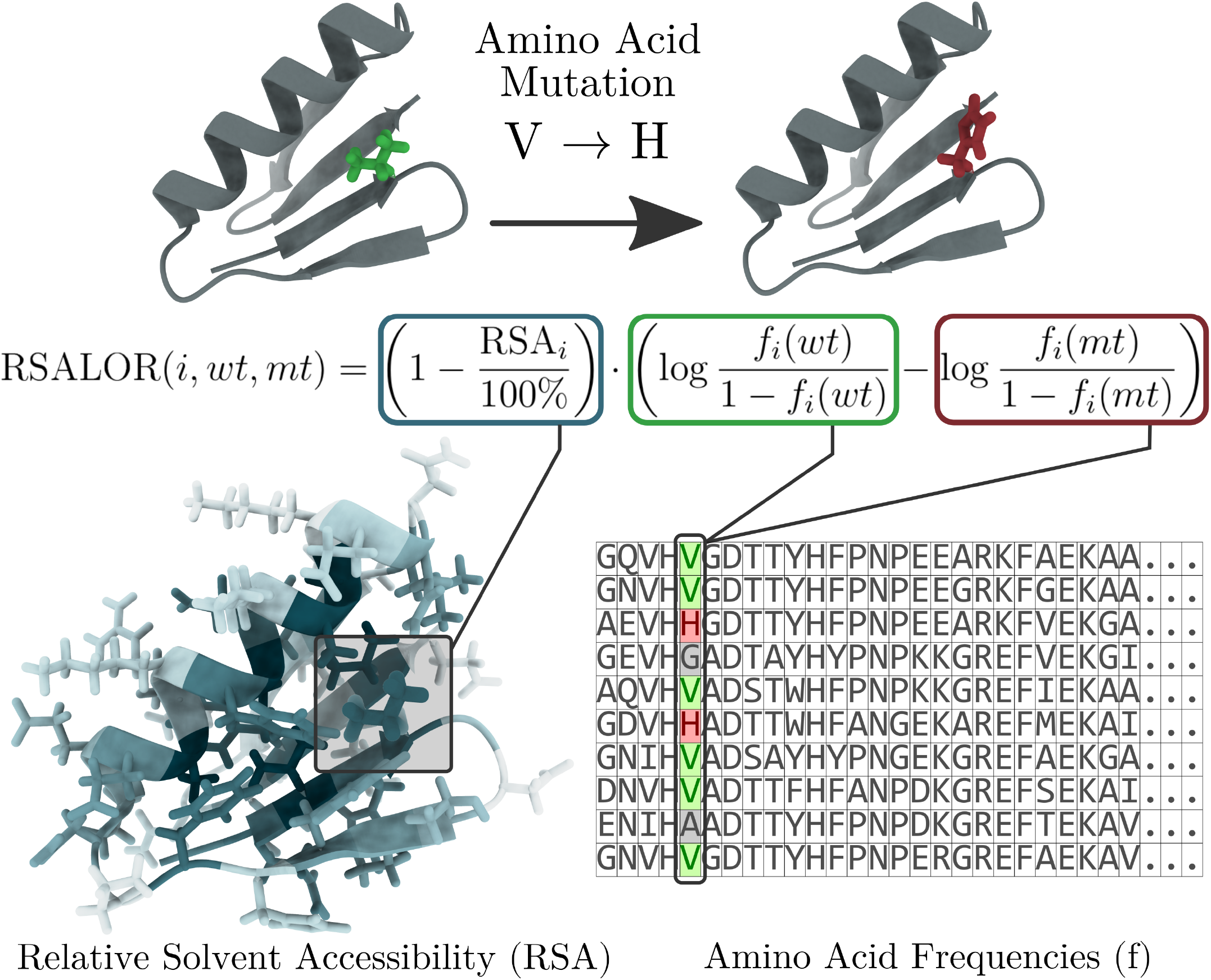
Graphical representation of the RSALOR model.

### RSALOR model

We present a very simple, independent-site, unsupervised approach for mutational effects prediction that combines evolutionary and structural information. In this section, we outline the key steps of the RSALOR model. Detailed implementation aspects are described in Supplementary Materials (SM) Sections 1 and 2.

The evolutionary information used in the model is derived from amino acid frequencies at the mutated position in a multiple sequence alignment (MSA), using the LOR between wild-type and mutant amino acid frequencies. The MSA is first curated by removing redundant identical sequences and those falling within the “twilight zone” [8] (i.e., sequences too evolutionary distant from the target sequence based on a sequence identity criterion). Indeed, we observed that the presence of very distant sequences adds noise rather than improving RSALOR predictions.

Since MSAs can be dominated by clusters of closely related sequences, we computed the “weighted” amino acid frequencies by reducing the contribution of sequences from larger clusters, as done in coevolutionary models, e.g. [9, 10]. We discussed and analyzed the impact of the weighting step in SM Section 3.2, showing that it mildly but consistently improves our model’s performance. To prevent LOR values from diverging and to handle the lack of information in small MSAs, we applied regularization to these frequencies. Using the weighted and regularized frequencies *f*_*i*_(*wt*) and *f*_*i*_(*mt*) for the wild-type and mutant amino acids at position *i*, the LOR is defined as:

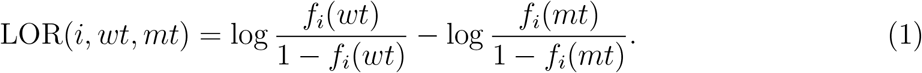

The sign of LOR is defined such that the result of mutations from a highly represented amino acid *wt* to a less represented amino acid *mt* is positive, which generally corresponds to a decrease in protein stability or fitness.

As structural information, we used the per-residue RSA, reflecting the observation that mutations in the protein core tend to have a greater impact than those on the surface. The simple product of LOR and the “complement” of RSA defines the RSALOR:

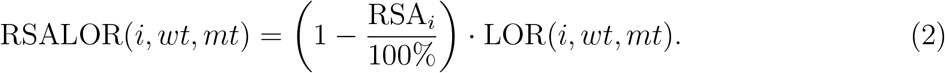

Since the RSA factor in this equation is always positive, the sign of RSALOR is the same as the sign of LOR.

Note that RSALOR nearly perfectly preserves the symmetry property (i.e., the effect of a mutation from *wt* to *mt* is opposite to the effect of the mutation from *mt* to *wt*). Indeed, while the evolutionary component LOR is perfectly symmetric, the fact that RSA is calculated using only the wild-type structure can, in principle, introduce asymmetry into the model. However, as shown in SM Section 3.4, this approximation does not substantially impact the predictions, which remain almost perfectly symmetric. This symmetry, which is often violated by predictors, has been shown to be an important feature in the prediction of changes in protein stability and binding affinity [11, 12, 13].

### RSALOR implementation

We provide RSALOR as a freely available, easy-to-install, and user-friendly Python package. It can be installed by cloning our GitHub repository at github.com/3BioCompBio/RSALOR or via the Python Package Index (PyPI) using **pip**. It takes as input the MSA of the target protein and its three-dimensional structure in PDB format.

The package automatically maps RSA values extracted from the structure to the corresponding positions in the MSA, even if the template structure contains missing residues or is homologous but not identical to the target sequence of the MSA. Indeed, RSA values are relatively robust to small structural changes. The model’s performance therefore remains almost unchanged when using structures with a few amino acid substitutions (see SM Section 3.5 for details).

The RSALOR package outputs or saves to a CSV file the following information for each possible single-site mutation in the target protein: the frequencies of gaps, wild-type and mutant residues in the MSA; the RSA of the mutated residue; and the LOR and RSALOR scores of the mutation.

### RSALOR performances

In [7], we evaluated RSALOR on its ability to predict the impact of mutations on protein stability. To further assess the robustness of the model, we tested it on ProteinGym [6] consisting of 218 standardized deep mutational scanning (DMS) experiments, covering a total of million mutations with annotated experimental effects on protein stability, binding affinity, fitness, activity, and expression.

**Table 1.**
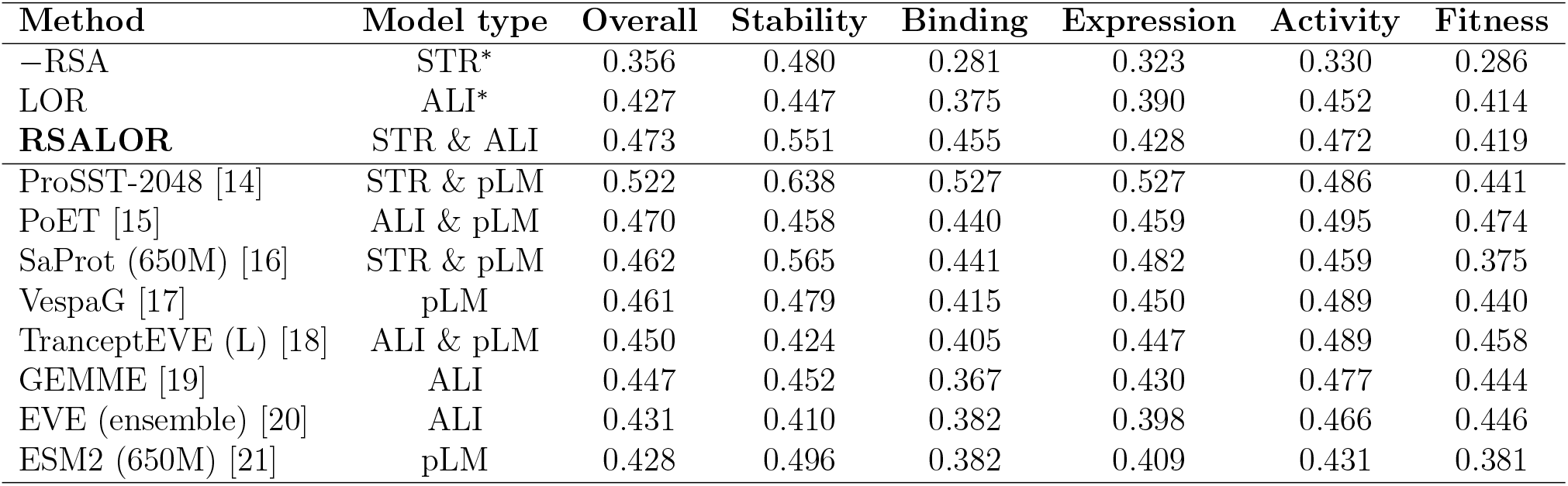
Average per-DMS Spearman correlations across ProteinGym subclasses (categorized by DMS target properties), comparing the RSALOR model with some of the top-performing models from the ProteinGym benchmark. Only single-site mutations were considered. Note that ProteinGym’s benchmark uses a slightly different method of averaging correlations, so their values and ours do not always perfectly match. ^*^STR = structure-based, ALI = alignment-based

To ensure a fair comparison with the other benchmarked models, we used the MSAs and structures provided by the ProteinGym repository without any modifications. These structures are AlphaFold-generated models [22], as the target sequences of most DMS experiments are not, or only partially, covered by experimental structures. Importantly, this means that our model does not rely on the availability of high-quality experimental structures. We additionally assessed the robustness of our predictions using alternative MSAs and structures, and observed essentially the same results (see details in SM Section 3.5).

While we present here a summary of the results, a more comprehensive analysis is provided in SM Section 3.1. It includes both overall performances and per-category performances on single-site and all (single-site and multiple) mutations from ProteinGym, evaluated using various metrics and compared among 27 different predictors (including 19 pLM-based models).

We first focused on the 700, 000 single-site mutations in the dataset. In Tab. 1, we present a comparison of the Spearman correlations, averaged over all DMS experiments or over specific DMS categories. The predictions of RSALOR were compared with some of the top-performing tools included in the ProteinGym benchmark. Our results show that the simple RSALOR model achieves performance in line with the other state-of-the-art methods, with an average Spearman correlation of 0.473 across all 218 datasets. Among the 27 models in the unsupervised category, only ProSST [14], which combines structure and pLM features, achieves better results. We note that the structural contribution, RSA, has a variable impact on performance depending on the DMS target property. Although it provides great improvements to the LOR score on stability and binding datasets, it enhances accuracy to a lesser extent for expression, activity, and fitness datasets. This is consistent considering that stability and binding affinity are more directly related to protein structure. For instance, the RSA score alone outperforms most benchmarked predictions on stability datasets.

In addition, we have shown that using RSA computed from more appropriate input 3D conformations can further boost predictions (see SM Section 3.6). For example, when studying the impact of mutations on protein–protein binding affinity, using the structure of the protein complex instead of the monomeric structure provided by the ProteinGym dataset leads to substantially improved performance.

It is worth noting, as already highlighted in the ProteinGym benchmark [6], that predictions also vary greatly between datasets, with some DMSs being exceptionally well predicted (Spearman correlation above 0.7), while in others, all methods essentially fail. In contrast, the number of homologous sequences in the input MSA has only a minor impact on the performance of RSALOR (see details in SM Section 3.3).

To predict the effect of multiple mutations using the RSALOR model, we made the approximation that there are no epistatic effects. Therefore, the effect of a multiple mutation is simply the sum of the effects of its individual single-site mutations. Even with this simplistic assumption, RSALOR achieved a correlation of 0.484 on all ProteinGym mutations, outperformed only by ProSST [14] and PoET [15] (with correlations of 0.523 and 0.490, respectively). All values are provided in SM Section 3.1.

We would like to underline that the RSALOR model is clearly a rough approximation for estimating the effects of protein mutations. First, it assumes that mutations at fully exposed residues (with a RSA of 100%) have no effect on the protein. While the link between the RSA of mutated residues and mutational impacts is well known [23, 24], the strength of its effect can vary significantly depending on the biophysical property considered. Second, RSALOR completely ignores epistatic effects and potential evolutionary information from other residue positions. While their contribution could be less significant in describing protein stability [7, 25], they seem to play an essential roles in protein activity and fitness [25, 26]. Despite these limitations, the model is on par with or slightly outperforms nearly all benchmarked models, highlighting that predicting mutational landscapes remains a challenge for current state-of-the-art methods.

Remarkably, combining the predictions of other models with RSA values (using Eq. 2) substantially improves the performance for almost all of the 27 benchmarked predictors. This holds true even for models that already incorporate structural information as input (see details in SM Section 4). We thus show that effectively incorporating RSA and other types of structural knowledge into evolutionary-or pLM-based models can lead to improved performance.

Finally, an important feature of RSALOR is its ease of use. It requires no model training or external dependencies, is easily installed with a single **pip** command, and runs in a straight-forward manner (see the GitHub repository). From a computational perspective, RSALOR is highly efficient. Its weighting step, being the most computationally intensive, is implemented in C++ and supports multi-threading. For instance, we evaluated the 2.5 millions mutations from ProteinGym in less than 20 minutes on a laptop using 8 CPUs. Results for each individual protein were generated in a time range of 1 second to 1 minute.

## Acknowledgments

We acknowledge financial support from the Belgian Fund for Scientific Research (F.R.S.-FNRS) through a PDR project. M.T. benefits from a FNRS-FRIA PhD grant. P.H. benefits from a Win4Doc grant from SPW Recherche of the Walloon Region.

