## Supplementary material for "Residue conservation and solvent accessibility are (almost) all you need for predicting mutational effects in proteins": Supp Material

**4 Combine RSA with other models** **19**

### 1 Installation and usage

#### 1.1 Installation and requirements

The RSALOR package can be installed easily using the Python standard package manager pip with the following command: `pip install rsalor`.

The source code of the package can be found in the GitHub repository `https://github.com/3BioCompBio/RSALOR`.

The package does not require external software dependencies and runs on Python version 3.9 or later. It depends on only two Python packages: Biopython (version 1.75 or later) (Cock et al., 2009) and NumPy (Harris et al., 2020). During installation, `pip` automatically compiles the C++ component of the package. This requires a C++ compiler (such as GCC) to be installed on the system.

Alternatively (without using pip), users can build the package directly from the source code. In this case, the C++ components must be compiled manually following the instructions provided in the repository's `README`.

#### 1.2 Usage

We have designed the `rsalor` package to be as easy to use as possible. The module requires three inputs: a path to an MSA file (in `.fasta` or `.a2m` format), a path to a three-dimensional (3D) structure file (in `.pdb` format), and the chain ID to consider in the protein structure. For example, in most computationally modeled monomeric structures, the only chain in the structure is typically named `"A"`.

> **Note:** Ensure that the first sequence in the MSA is the target protein sequence to be mutated. In addition, confirm that all sequences in the MSA file have the same length; otherwise, it is a list of sequences and not a valid MSA.

Suppose you have two files in your working directory: an MSA file `"./protein1.fasta"` and a 3D structure file `"./protein1.pdb"` with a chain ID `"A"`. You can compute the predicted mutational landscape of the protein using the `rsalor` package in just three lines of code:

```
from rsalor import MSA
msa = MSA("./protein1.fasta", "./protein1.pdb", "A", verbose=True)
rsalor_scores = msa.get_scores()
```

The output object `rsalor_scores` is a list of dictionaries containing the computed scores (LOR, RSALOR, and others), each assigned to a single-site mutation in the target protein. Mutations' notation is explained in Sections 2.1 and 2.7. How the different scores are computed is explained in Sections 2.5, 2.8 and 2.9.

Alternatively, you can directly save the computed scores to a `.csv` file (using the `sep` argument to specify the CSV separator):

```
msa.save_scores("./protein1_rsalor_scores.csv", sep=";")
```

Note that if you omit the path to the structure file and the chain, the package will still execute and compute the LOR and LR values without incorporating RSA information into its predictions:

```
from rsalor import MSA
msa = MSA("./protein1.fasta", verbose=True)
rsalor_scores = msa.get_scores() # scores without the RSA
```

It is also possible to save to files the RSA values computed on the protein structure, the weights assigned to each sequence of the MSA, and the filtered and trimmed MSA (as described in Sections 2.1 and 2.2) by specifying the following arguments:

```
msa = MSA(
    "./protein1.fasta", "./protein1.pdb", "A",
    weights_cache_path="./protein1_weights.txt",
    rsa_cache_path="./protein1_rsa.txt",
    trimmed_msa_path="protein1_msa_trimmed.fasta",
)
```

If you want to use the package to compute (and save) weights on the unfiltered MSA (exactly as in the input file), you can deactivate the filtering of the MSA by setting the optional arguments `remove_redundant_sequences=False` and `min_seqid=None`.

##### 1.3 Optional arguments

The `rsalor` package provides many optional arguments to customize the computation of its scores. These optional arguments can be specified during the initialization of the `MSA` object. You can view the complete list of optional arguments by running:

```
from rsalor import MSA
MSA.help()
```

- **theta_regularization** (`float`, default: `0.01`):
  Regularization term for LOR/LR at the amino acid frequencies level (see Section 2.4).
- **n_regularization** (`float`, default: `0.0`):
  Regularization term for LOR/LR at the amino acid counts level (see Section 2.4).
- **count_target_sequence** (`bool`, default: `True`):
  Inclusion of the target (first) sequence of the MSA when calculating frequencies.
- **remove_redundant_sequences** (`bool`, default: `True`):
  Preprocessing of the MSA to remove redundant sequences (see Section 2.2).
- **seqid_weights** (`None | float`, default: `0.80`):
  Sequence identity threshold for clustering sequences during weighting (see Section 2.3). Set to `None` to ignore weights.
- **min_seqid** (`None | float`, default: `0.35`):
  Discard sequences with a sequence identity to target below this threshold (see Section 2.2). Set to `None` to disable this filter.
- **num_threads** (`int`, default: `1`):
  Number of threads (CPUs) to use for weight evaluation in the C++ backend.
- **rsa_solver** (`"biopython"/"DSSP"/"MuSiC"`, default `"biopython"`):
  Solver used to compute RSA. Note that DSSP and MuSiC require their respective software to be installed.
- **rsa_solver_path** (`None | str`, default: `None`):
  Path to the DSSP or MuSiC executable for RSA computation. Leave as `None` if the software is in the system `PATH`.
- **trimmed_msa_path** (`None | str`, default: `None`):
  Path to save the trimmed and filtered MSA file (see Sections 2.1 and 2.2). Leave as `None` to ignore.

- **allow_msa_overwrite** (`bool`, default: `False`):
  Allow overwriting the initial MSA file with the trimmed and filtered version (if `msa_path` is the same as `trimmed_msa_path`).

- **weights_cache_path** (`None | str`, default: `None`):
  Path to read pre-computed weights (if the file exists) or save calculated weights (if the file does not exist). Leave empty to ignore.

- **rsa_cache_path** (`None | str`, default: `None`):
  Path to read pre-computed RSA values (if the file exists) or save calculated RSA values (if the file does not exist). Leave empty to ignore.

- **verbose** (`bool`, default: `False`):
  Enable logging of execution steps.

- **disable_warnings** (`bool`, default: `False`):
  Prevents warnings from being logged. By default, warnings are logged even if `verbose=False`.

- **name** (`None | str`, default: `None`):
  Identifier for the MSA object (used in logs).

### 2 Implementation details

The software RSALOR is available as a Python `pip` package (`rsalor`) and is implemented in Python and C++. Additionally, the code can also be used as a standalone python module. The weighting step is the most computationally intensive, and for that reason, it is implemented in C++ and supports multi-threading. The C++ component is seamlessly integrated into the Python code, remaining completely "invisible" to the user, and is automatically compiled during installation via `pip`.

The package takes as input an MSA and a protein 3D structure (along with the corresponding `chain_id` in the structure). It processes the MSA by parsing, filtering, and weighting its sequences (Sections 2.1, 2.2 and 2.3); and computes evolutionary mutational scores, LR and LOR (Sections 2.4 and 2.5). On the other hand, the package extracts the sequence from the structure file and calculates RSA values for each residue (Section 2.6). It then maps the RSA values to positions in the MSA (Section 2.7) and combines these values with the evolutionary scores to compute RSALR and RSALOR scores (Section 2.8). Finally, the software outputs (or saves in a `.csv` file) all the calculated scores for each single-site mutation in the target protein sequence (detailed in Section 2.9).

The package requires no external software dependencies to run. It relies on the Python library Biopython (Cock et al., 2009) to compute RSA values and to perform pairwise alignments between the target sequence and the sequence extracted from the structure file. To speed up the evaluation of evolutionary scores (such as calculation of frequencies and regularization), the package exploits array programming using the Python package NumPy (Harris et al., 2020).

#### 2.1 MSA trimming

It is common for an MSA generated from a reference sequence dataset to contain gapped positions in the target sequence. These positions are excluded when computing the effects of mutations in the protein and are therefore trimmed (i.e., removed from all sequences in the MSA). This trimming creates two references for naming mutations: one based on the initial input MSA (referred as `mutation_fasta`) and another based on the trimmed MSA (referred as `mutation_fasta_trimmed`). For simplicity, the output of the `rsalor` package always specifies both conventions, even if no trimming is done.

##### 2.2 Sequence filtering

The package first removes duplicate sequences (exact matches between two sequences in the MSA). Note that it is possible for two reference sequences to be initially different (as represented in the reference sequence database used to construct the MSA), but to have identical alignments in the trimmed MSA when only positions aligned to the target sequence of the MSA are considered. The impact of this filter on the resulting LOR and LR scores is marginal, since the sequences are weighted later. Indeed, keeping multiple copies of the same sequence in the MSA would simply reduce their individual weights. However, this filter can significantly improve the computation time of the weighting step, especially if the MSA contains many duplicates (which is often the case for MSAs of short sequences).

The package then removes all sequences from the "twilight zone" (Rost, 1999), i.e., sequences that are too distant from the target sequence based on a sequence identity criterion. While these sequences may be useful for more complex epistatic models, their mutational landscapes are too divergent from that of the target sequence to provide meaningful information in the context of an independent-site model. To avoid removing sequences that align well with the target sequence but only over a small fraction of the MSA positions (for instance, when the target sequence contains multiple domains and the aligned sequence contains only one domain), we compute sequence identity exclusively on aligned positions (i.e., positions that are not gaps in the aligned sequence).

By default, the sequence identity threshold is set to 0.35, following (Rost, 1999). Both the duplicate filter and the twilight zone filter can be disabled.

##### 2.3 Sequence weighting

Similarly to Direct Coupling Analysis (DCA) methods (Cocco et al., 2018), the `rsalor` package relatively weights sequences of the input MSA. Since MSAs often contain clusters of very closely related proteins, these weights help mitigate bias toward overrepresented clusters of sequences.

For an MSA containing $N$ sequences, where the set of sequences is denoted as $S$, the weight $w_j$ of a sequence $s_j$ is defined as the inverse of $m_j$, the number of sequences in the MSA sharing a sequence identity with $s_j$ greater than or equal to a threshold value $\tau$:

$$w_j = \frac{1}{m_j} = \frac{1}{|\{s \in S \mid \mathrm{seqid}(s_j, s) \geq \tau\}|}. \quad (1)$$

Once the sequences are weighted, we define $N_{\mathrm{eff}}$, the effective number of sequences in the MSA as:

$$N_{\mathrm{eff}} = \sum_{j=1}^{N} w_j. \quad (2)$$

Rather than using the regular frequency $f_i(a)$ of an amino acid type $a$ at position $i$ defined as:

$$f_i(a) = \frac{\sum_{j=1}^{N} \delta(\mathrm{seq}_j(i), a)}{N}, \quad (3)$$

we use the weighted frequency $f_i^w(a)$ of an amino acid type $a$ at position $i$ as:

$$f_i^w(a) = \frac{\sum_{j=1}^{N} \delta(\mathrm{seq}_j(i), a) \cdot w_j}{N_{\mathrm{eff}}}, \quad (4)$$

where $\mathrm{seq}_j(i)$ represents the amino acid type at position $i$ of the $j^{\mathrm{th}}$ sequence and $\delta$ is the Kronecker delta function.

By default, the sequence identity threshold $\tau$ is set to 0.80 (as usually done in DCA methods (Pucci et al., 2024) and in our previous work (Hermans et al., 2024)). The weighting step can be omitted, which significantly accelerates execution. However, weights have been shown to improve prediction accuracy (Hermans et al., 2024), and the overall weighting computation is relatively fast (taking at most one minute on a laptop using 8 CPUs for any protein from the ProteinGym deep mutational scanning (DMS) dataset collection (Notin et al., 2024)).

The weighting step involves performing an all-to-all comparison of all sequences in the MSA, making it the most computationally intensive part of the package. For performance reasons, this step is implemented in C++ and supports multi-threading. The integration between Python and C++ is partly inspired by the pip package `pycofitness` implementation (Pucci et al., 2024).

For the sake of simplicity, we will omit to specify that all frequencies are weighted in the following equations and text.

##### 2.4 Frequency regularization

If a frequency $f_i(a)$ is equal to zero (amino acid type $a$ is never observed at position $i$ in the MSA) or one (only amino acid type $a$ is observed at position $i$ in the MSA), the resulting LOR and LR values would diverge due to the logarithmic calculations they involve. To address this issue and to handle the lack of information in small MSAs, it is necessary to regularize the observed frequencies in the MSA.

The package implements two types of regularization: frequency-based and count-based. The main difference between them is that the first regularization remains constant as the MSA depth increases, whereas the latter becomes less prevalent.

Considering an MSA of length $N$, the frequency $f_i(a)$ is computed as $f_i(a) = \frac{c_i(a)}{N}$, where $c_i(a)$ is the count of amino acids of type $a$ at position $i$. We define the count-based regularization with amplitude $n$ as:

$$\tilde{f_i}^n(a) = \frac{c_i(a) + n}{N + 21n}. \tag{5}$$

Note that 21 represents the number of possible states (20 amino acid types and one gap). On the other hand, we define frequency-based regularization of amplitude $\theta$ as,

$$\tilde{f_i}^\theta(a) = f_i(a)(1 - \theta) + \frac{\theta}{21}. \tag{6}$$

When both regularization methods are applied, the count-based regularization is applied before the frequency-based one, resulting in the following equation:

$$\tilde{f_i}^{n,\theta}(a) = \frac{(c_i(a) + n)(1 - \theta)}{N + 21n} + \frac{\theta}{21}. \tag{7}$$

For the regularization of weighted frequencies, the same equation holds, with $N$ replaced by $N_{\text{eff}}$ and $c_i(a)$ replaced by its weighted counterpart, $\sum_{j=1}^{N} \delta(\text{seq}_j(i), a) \cdot w_j$.

By default, the `rsalor` package only uses frequency-based regularization with $\theta = 0.01$, as we did previously in (Hermans et al., 2024) and (Tsishyn et al., 2024a). However, we allow the user to incorporate count-based regularization, as it is also a widely used regularization method in models predicting mutation effects such as GEMME (Laine et al., 2019). Note that we do not recommend using only count-based regularization, as it can lead to very extreme LOR values for deep MSAs.

For the sake of simplicity, we will omit specifying that all frequencies are regularized in the following equations and text.

##### 2.5 Evolutionary mutational scores

The package assigns an "evolutionary energy" score to each possible single-site mutation in the target protein. We define the Log-Odd Ratio (LOR) (Raimondi et al., 2016) and the Log Ratio (LR) of a mutation at position $i$ from the wild-type amino acid $wt$ to the mutant amino acid $mt$ as:

$$\mathrm{LOR}(i, wt, mt) = \log \frac{f_i(wt)}{1 - f_i(wt)} - \log \frac{f_i(mt)}{1 - f_i(mt)}, \tag{8}$$

$$\mathrm{LR}(i, wt, mt) = \log f_i(wt) - \log f_i(mt). \tag{9}$$

The LOR and LR values are computed using the weighted and regularized frequencies. The sign of LOR (the same applies to LR) is defined such that a mutation from a highly represented amino acid $wt$ to a poorly represented amino acid $mt$ results in a positive LOR, which generally corresponds to a disruptive mutation.

While the LR score naturally arises from the Boltzmann distribution as an estimate of the "evolutionary energy", we noticed in our previous works (Hermans et al., 2024; Tsishyn et al., 2024a) that the LOR score is consistently slightly more predictive than the LR score (at least when used alone or combined with the RSA). Nevertheless, the LR score is retained in our package as it may be relevant as a feature and could be more suitable for specific applications. For instance, the LR score corresponds to the optimal solution for the restriction to independent-site models in DCA using the maximum entropy principle (Cocco et al., 2018).

Note also that since LOR (the same applies to LR) is a state function, its value can be computed between any two amino acid types $a_1$ and $a_2$ (not exclusively starting from the wild-type amino acid).

#### 2.6 Relative solvent accessibility evaluation

The package can compute the Relative Solvent Accessibility (RSA) using two different solvers: Biopython module `Bio.PDB.SASA` (Cock et al., 2009) and the widely used software DSSP (Kabsch and Sander, 1983). While the DSSP solver requires the DSSP software to be installed on the machine, the Biopython solver runs without any external dependencies. For that reason, Biopython is set as the default solver of the `rsalor` package.

The two solvers are based on the Shrake and Rupley algorithm (Shrake and Rupley, 1973), which samples spheres of a given radius (simulating molecules of the solvent) to "probe" the surface of the molecule. The algorithm computes the solvent accessible surface area (SASA), which is then normalized to RSA (in %) by dividing the SASA of the residue in its structure by the SASA of the same residue in an extended tripeptide Gly-X-Gly conformation (Rose et al., 1985). For the Biopython solver, we use the empirically computed scale from (Tien et al., 2013) to convert SASA to RSA values.

While the main algorithm is the same for the two solvers, some implementation differences lead to slightly different RSA values. However, these differences have almost no impact on the predictions, with an average Spearman correlation of 0.995 between the RSALOR values obtained by the two solvers on the single-site mutations from ProteinGym.

#### 2.7 Align sequence and structure

It is common that the target sequence and the sequence extracted from the protein 3D structure do not perfectly match. For instance, most experimental protein structures from the Protein Data Bank (PDB) (Berman et al., 2000) contain missing residues (especially at the termini of the protein chain) or non-standard amino acids. Additionally, residue indices in the PDB 3D structure often do not correspond to sequential indices assigned in a FASTA sequence. Another example of such a mismatch is when the provided MSA and the 3D structure represent slightly different segments of a larger protein (as is the case for some proteins in ProteinGym (Notin et al., 2024)). Finally, a user might wish to evaluate the mutational landscape of a protein using the RSALOR score derived from the structure of a homologous template protein that differs slightly from the target protein, assuming the overall secondary and tertiary structures are conserved. This scenario is reasonable, as the RSA value of a residue tends to remain similar even if it is mutated.

All these scenarios make the manual process of mapping RSA values extracted from the structure to positions in the MSA tedious. The package automates this process by performing pairwise alignments between the target sequence and the sequence extracted from the protein 3D structure. After extracting the sequence from the structure file, it converts any non-standard amino acids in the protein structure to their corresponding standard amino acids and then performs pairwise alignments using the Biopython module `Bio.Align.PairwiseAligner`.

This alignment process gives rise to a third reference for naming mutations (in addition to the two defined in Section 2.1): one based on the mutated residue as it is referenced in the structure file (referred to as `mutation_pdb`). The output of the `rsalor` package also specifies this mutation naming convention.

##### 2.8 Combine RSA with evolutionary scores

Finally, the package combines the evolutionary score LOR (and LR) with the RSA of the mutated residue. The RSA value is used to modulate the amplitude of the evolutionary score. To reflect the anti-correlation relationship (Hermans et al., 2024) between the RSA of a residue and the average amplitude of the effect of mutations at this residue, we calculate the complement of RSA and rescale it between zero and one. Furthermore, to ensure that the RSA-based factor does not invert the sign of the evolutionary score, we cap RSA values at 100% (to handle cases where RSA exceeds 100%). This gives us a score that can be described as a "relative solvent inaccessibility ratio". The final scores, RSALOR and RSALR, are defined as follows:

$$\mathrm{RSALOR}(i, wt, mt) = \left( \frac{100\% - \min\{\mathrm{RSA}_i, 100\%\}}{100\%} \right) \cdot \mathrm{LOR}(i, wt, mt), \tag{10}$$

$$\mathrm{RSALR}(i, wt, mt) = \left( \frac{100\% - \min\{\mathrm{RSA}_i, 100\%\}}{100\%} \right) \cdot \mathrm{LR}(i, wt, mt), \tag{11}$$

where $\mathrm{RSA}_i$ is the RSA of the residue at position $i$.

##### 2.9 Final output

Starting from an MSA and a protein 3D structure, the module performs computational DMS by evaluating each possible single-site mutation of the target protein sequence. For each mutation, it provides the following: the mutation represented using the three naming conventions (`mutation_fasta`, `mutation_fasta_trimmed` and `mutation_pdb`); the weighted but not regularized frequencies at this position of gaps, wild-type amino acids and mutant amino acids; the RSA of the mutated residue; and the mutational scores LR, LOR, RSALR and RSALOR.

Since the model considers the different sites as independent by design, it does not provide predictions for multiple mutations. Indeed, the model does not contain any information about how the effects of two or more mutations might interact. To address this for predicting the effects of multiple mutations in ProteinGym, we assume the absence of epistatic effects and estimate the combined impact by summing the effects of each individual mutation. Surprisingly, this led to results that are in line with the best epistatic methods.

### 3 Performance and robustness of RSALOR

#### 3.1 Global performances on ProteinGym

In the main paper, we compared the Spearman correlations of LOR, RSALOR and a selection of other prediction models with single-site DMS scores from ProteinGym. Here, we provide a comprehensive analysis that includes all the models presented in ProteinGym (accessed March 2025) on single-site and multiple mutations, using both Spearman correlation and additional metrics.

ProteinGym currently benchmarks 71 models (in the zero-shot, DMS substitution category). However, many are variations of the same model. For instance, protein language models are typically represented by multiple versions, each containing a different number of parameters. We retained only the best-performing model in each publication (based on ProteinGym's global ranking). This gave us 27 distinct predictors representing a wide range of approaches, including many multiple sequence alignment-based models (ALI), protein language models (pLM), and two inverse folding models (IFm). Some of these models also exploit structural information (STR). Several hybrid models combine two of the aforementioned approaches, as is the case for RSALOR that combines sequence alignment and structural information.

Since we do not expect a linear relationship between predicted scores and DMS scores (Boucher et al., 2016), the Spearman correlation is the most suitable metric to evaluate performance in our case (Notin et al., 2024). We present Spearman correlations for LOR, RSALOR and the 27 benchmarked models on single-site mutations in Tab. S1 and on all mutations in Tab. S2. Results are reported as the average of the Spearman correlations computed on each DMS dataset and are also categorized according to the properties targeted by the DMS, i.e., stability, binding, expression, activity, and fitness.

Despite its simplicity and independent-site nature, RSALOR ranks second and third for single-site and all mutations, respectively. We graphically represent in Fig. S1 and S2 the Spearman correlations of RSALOR and the 27 benchmarked predictors on single-site and all mutations, both on all DMS datasets and within the specific DMS categories considered. We observe that RSALOR performs relatively well on all DMS categories, with the highest score reached on stability.

We also reported the predictors' performances measured by the Pearson correlation, the Matthews correlation coefficient (MCC), and the area under the receiver operating characteristic curve (AUC) for both single-site and all mutations in Tab. S3. Although the Pearson correlation assumes a linear relationship between compared values, it remains widely used and supports observations from Spearman correlation. In contrast, AUC and MCC metrics compare predicted scores with binarized experimental DMS scores, providing complementary information to rank-based correlations, especially when DMS measurements exhibit a bimodal distribution. RSALOR consistently shows good performances with all these metrics.

One final note: ProteinGym's benchmark uses a slightly different method for averaging correlations, so their values and ours do not always match perfectly. However, these differences are small and do not affect the ranking of RSALOR.

**Table S1: Spearman correlations between predicted values and experimental DMS scores for single-site mutations**. Average per-DMS Spearman correlations for all data (column 3) and categorized by target properties (columns 4-8), for LOR, RSALOR and the 27 benchmarked models. For references to the prediction methods, see ProteinGym repository (Notin et al., 2024).

| Method | Model type | Overall | Stability | Binding | Expression | Activity | Fitness |
|---|---|---|---|---|---|---|---|
| LOR | ALI | 0.427 | 0.447 | 0.375 | 0.390 | 0.452 | 0.414 |
| **RSALOR** | STR & ALI | 0.473 | 0.551 | 0.455 | 0.428 | 0.472 | 0.419 |
| ProSST (2048) | STR & pLM | 0.522 | 0.638 | 0.527 | 0.527 | 0.486 | 0.441 |
| PoET | ALI & pLM | 0.470 | 0.458 | 0.440 | 0.459 | 0.495 | 0.474 |
| ProtSSN (ens) | STR & pLM | 0.464 | 0.541 | 0.446 | 0.442 | 0.458 | 0.410 |
| SaProt (650M) | STR & pLM | 0.462 | 0.565 | 0.441 | 0.482 | 0.459 | 0.375 |
| VespaG | pLM | 0.461 | 0.479 | 0.415 | 0.450 | 0.489 | 0.440 |
| TranceptEVE (L) | ALI & pLM | 0.450 | 0.424 | 0.405 | 0.447 | 0.489 | 0.458 |
| GEMME | ALI | 0.447 | 0.452 | 0.367 | 0.430 | 0.477 | 0.444 |
| VESPA | pLM | 0.442 | 0.439 | 0.440 | 0.400 | 0.466 | 0.442 |
| TranceptEVE (S) | ALI & pLM | 0.439 | 0.417 | 0.397 | 0.433 | 0.476 | 0.446 |
| ESM-IF1 | IFm | 0.434 | 0.624 | 0.446 | 0.400 | 0.353 | 0.323 |
| MIFST | STR & pLM | 0.432 | 0.524 | 0.399 | 0.431 | 0.404 | 0.375 |
| EVE (ens) | ALI | 0.431 | 0.410 | 0.382 | 0.398 | 0.466 | 0.446 |
| MSA-Transformer (ens) | ALI & pLM | 0.430 | 0.429 | 0.322 | 0.438 | 0.470 | 0.425 |
| ESM2 (650M) | pLM | 0.428 | 0.496 | 0.382 | 0.409 | 0.431 | 0.381 |
| ESM1v (ens) | pLM | 0.409 | 0.423 | 0.359 | 0.421 | 0.419 | 0.398 |
| DeepSequence (ens) | ALI | 0.407 | 0.393 | 0.361 | 0.379 | 0.449 | 0.411 |
| CARP (640M) | pLM | 0.396 | 0.425 | 0.304 | 0.398 | 0.407 | 0.379 |
| ESM1b | pLM | 0.389 | 0.428 | 0.340 | 0.396 | 0.401 | 0.356 |
| Progen2 (XL) | pLM | 0.386 | 0.381 | 0.365 | 0.408 | 0.395 | 0.385 |
| EVmutation | ALI | 0.381 | 0.321 | 0.329 | 0.372 | 0.437 | 0.413 |
| RITA (XL) | pLM | 0.360 | 0.332 | 0.336 | 0.406 | 0.363 | 0.376 |
| Wavenet | ALI | 0.358 | 0.367 | 0.349 | 0.342 | 0.350 | 0.362 |
| Site-Independent | ALI | 0.340 | 0.278 | 0.340 | 0.341 | 0.370 | 0.377 |
| Unirep (evotuned) | ALI & pLM | 0.322 | 0.274 | 0.319 | 0.363 | 0.352 | 0.337 |
| MULAN (S) | pLM | 0.292 | 0.337 | 0.359 | 0.378 | 0.285 | 0.226 |
| ProteinMPNN | IFm | 0.282 | 0.550 | 0.165 | 0.194 | 0.179 | 0.149 |
| ProtGPT2 | pLM | 0.181 | 0.202 | 0.180 | 0.187 | 0.182 | 0.161 |

**Table S2: Spearman correlations between predicted values and experimental DMS scores for all (single-site and multiple) mutations**. Average per-DMS Spearman correlations for all data (column 3) and categorized by target properties (columns 4-8), for LOR, RSALOR and the 27 benchmarked models. For references to the prediction methods, see ProteinGym repository (Notin et al., 2024).

| Method | Model type | Overall | Stability | Binding | Expression | Activity | Fitness |
|---|---|---|---|---|---|---|---|
| LOR | ALI | 0.452 | 0.508 | 0.372 | 0.395 | 0.463 | 0.426 |
| **RSALOR** | STR & ALI | 0.484 | 0.574 | 0.412 | 0.432 | 0.483 | 0.433 |
| ProSST (2048) | STR & pLM | 0.523 | 0.653 | 0.449 | 0.530 | 0.486 | 0.442 |
| PoET | ALI & pLM | 0.490 | 0.519 | 0.384 | 0.466 | 0.501 | 0.482 |
| ProtSSN (ens) | STR & pLM | 0.473 | 0.568 | 0.359 | 0.449 | 0.472 | 0.416 |
| SaProt (650M) | STR & pLM | 0.473 | 0.592 | 0.371 | 0.488 | 0.466 | 0.388 |
| VespaG | pLM | 0.482 | 0.533 | 0.363 | 0.456 | 0.498 | 0.455 |
| TranceptEVE (L) | ALI & pLM | 0.474 | 0.500 | 0.359 | 0.457 | 0.492 | 0.466 |
| GEMME | ALI | 0.474 | 0.519 | 0.368 | 0.438 | 0.485 | 0.456 |
| VESPA | pLM | 0.463 | 0.500 | 0.362 | 0.404 | 0.478 | 0.454 |
| TranceptEVE (S) | ALI & pLM | 0.468 | 0.497 | 0.376 | 0.443 | 0.480 | 0.458 |
| ESM-IF1 | IFm | 0.437 | 0.624 | 0.383 | 0.407 | 0.359 | 0.335 |
| MIFST | STR & pLM | 0.415 | 0.485 | 0.309 | 0.438 | 0.393 | 0.379 |
| EVE (ens) | ALI | 0.460 | 0.491 | 0.369 | 0.408 | 0.470 | 0.455 |
| MSA-Transformer (ens) | ALI & pLM | 0.453 | 0.492 | 0.317 | 0.446 | 0.479 | 0.428 |
| ESM2 (650M) | pLM | 0.438 | 0.523 | 0.327 | 0.415 | 0.436 | 0.391 |
| ESM1v (ens) | pLM | 0.428 | 0.477 | 0.305 | 0.429 | 0.425 | 0.408 |
| DeepSequence (ens) | ALI | 0.438 | 0.476 | 0.337 | 0.390 | 0.459 | 0.423 |
| CARP (640M) | pLM | 0.390 | 0.412 | 0.259 | 0.397 | 0.404 | 0.382 |
| ESM1b | pLM | 0.420 | 0.500 | 0.281 | 0.406 | 0.436 | 0.368 |
| Progen2 (XL) | pLM | 0.406 | 0.445 | 0.291 | 0.418 | 0.404 | 0.389 |
| EVmutation | ALI | 0.418 | 0.430 | 0.304 | 0.378 | 0.446 | 0.421 |
| RITA (XL) | pLM | 0.384 | 0.398 | 0.286 | 0.414 | 0.368 | 0.390 |
| Wavenet | ALI | 0.390 | 0.449 | 0.314 | 0.350 | 0.379 | 0.367 |
| Site-Independent | ALI | 0.369 | 0.358 | 0.324 | 0.343 | 0.378 | 0.386 |
| Unirep (evotuned) | ALI & pLM | 0.353 | 0.366 | 0.280 | 0.365 | 0.362 | 0.347 |
| MULAN (S) | pLM | 0.304 | 0.369 | 0.320 | 0.383 | 0.286 | 0.237 |
| ProteinMPNN | IFm | 0.295 | 0.565 | 0.160 | 0.198 | 0.194 | 0.166 |
| ProtGPT2 | pLM | 0.198 | 0.257 | 0.139 | 0.193 | 0.183 | 0.166 |

**Table S3: Different performance metrics between predicted scores and experimental DMS scores for single-site and all mutations**. Average per-DMS Pearson correlations ($r$), AUC values and MCC values on single-site (-single) and all (-all) mutations for LOR, RSALOR and the 27 benchmarked models. For references to the prediction methods, see ProteinGym repository (Notin et al., 2024).

| **Method** | **Model type** | $r$-single | $r$-all | AUC-single | AUC-all | MCC-single | MCC-all |
|---|---|---|---|---|---|---|---|
| LOR | ALI | 0.428 | 0.448 | 0.741 | 0.751 | 0.336 | 0.359 |
| **RSALOR** | STR & ALI | 0.488 | 0.486 | 0.768 | 0.768 | 0.372 | 0.386 |
| ProSST (2048) | STR & pLM | 0.563 | 0.544 | 0.794 | 0.788 | 0.403 | 0.414 |
| PoET | ALI & pLM | 0.481 | 0.494 | 0.768 | 0.773 | 0.366 | 0.387 |
| ProtSSN (ens) | STR & pLM | 0.469 | 0.471 | 0.759 | 0.763 | 0.359 | 0.377 |
| SaProt (650M) | STR & pLM | 0.472 | 0.478 | 0.759 | 0.761 | 0.361 | 0.374 |
| VespaG | pLM | 0.394 | 0.403 | 0.765 | 0.770 | 0.362 | 0.383 |
| TranceptEVE (L) | ALI & pLM | 0.442 | 0.464 | 0.754 | 0.764 | 0.354 | 0.375 |
| GEMME | ALI | 0.453 | 0.471 | 0.755 | 0.763 | 0.351 | 0.373 |
| VESPA | pLM | 0.441 | 0.454 | 0.756 | 0.762 | 0.356 | 0.374 |
| TranceptEVE (S) | ALI & pLM | 0.430 | 0.457 | 0.748 | 0.760 | 0.347 | 0.369 |
| ESM-IF1 | IFm | 0.440 | 0.438 | 0.739 | 0.737 | 0.332 | 0.343 |
| MIFST | STR & pLM | 0.426 | 0.411 | 0.738 | 0.726 | 0.330 | 0.320 |
| EVE (ens) | ALI | 0.418 | 0.448 | 0.744 | 0.756 | 0.344 | 0.364 |
| MSA-Transformer (ens) | ALI & pLM | 0.429 | 0.447 | 0.742 | 0.751 | 0.336 | 0.359 |
| ESM2 (650M) | pLM | 0.441 | 0.444 | 0.742 | 0.744 | 0.336 | 0.347 |
| ESM1v (ens) | pLM | 0.416 | 0.429 | 0.729 | 0.737 | 0.319 | 0.339 |
| DeepSequence (ens) | ALI | 0.399 | 0.428 | 0.730 | 0.744 | 0.326 | 0.347 |
| CARP (640M) | pLM | 0.403 | 0.396 | 0.723 | 0.714 | 0.308 | 0.302 |
| ESM1b | pLM | 0.396 | 0.423 | 0.719 | 0.735 | 0.303 | 0.333 |
| Progen2 (XL) | pLM | 0.372 | 0.386 | 0.720 | 0.729 | 0.300 | 0.321 |
| EVmutation | ALI | 0.369 | 0.408 | 0.715 | 0.731 | 0.306 | 0.329 |
| RITA (XL) | pLM | 0.342 | 0.360 | 0.703 | 0.716 | 0.281 | 0.305 |
| Wavenet | ALI | 0.358 | 0.384 | 0.705 | 0.720 | 0.280 | 0.310 |
| Site-Independent | ALI | 0.310 | 0.346 | 0.694 | 0.704 | 0.279 | 0.297 |
| Unirep (evotuned) | ALI & pLM | 0.302 | 0.326 | 0.687 | 0.700 | 0.253 | 0.283 |
| MULAN (S) | pLM | 0.307 | 0.315 | 0.661 | 0.665 | 0.221 | 0.233 |
| ProteinMPNN | IFm | 0.272 | 0.290 | 0.651 | 0.660 | 0.209 | 0.227 |
| ProtGPT2 | pLM | 0.172 | 0.188 | 0.602 | 0.609 | 0.145 | 0.159 |

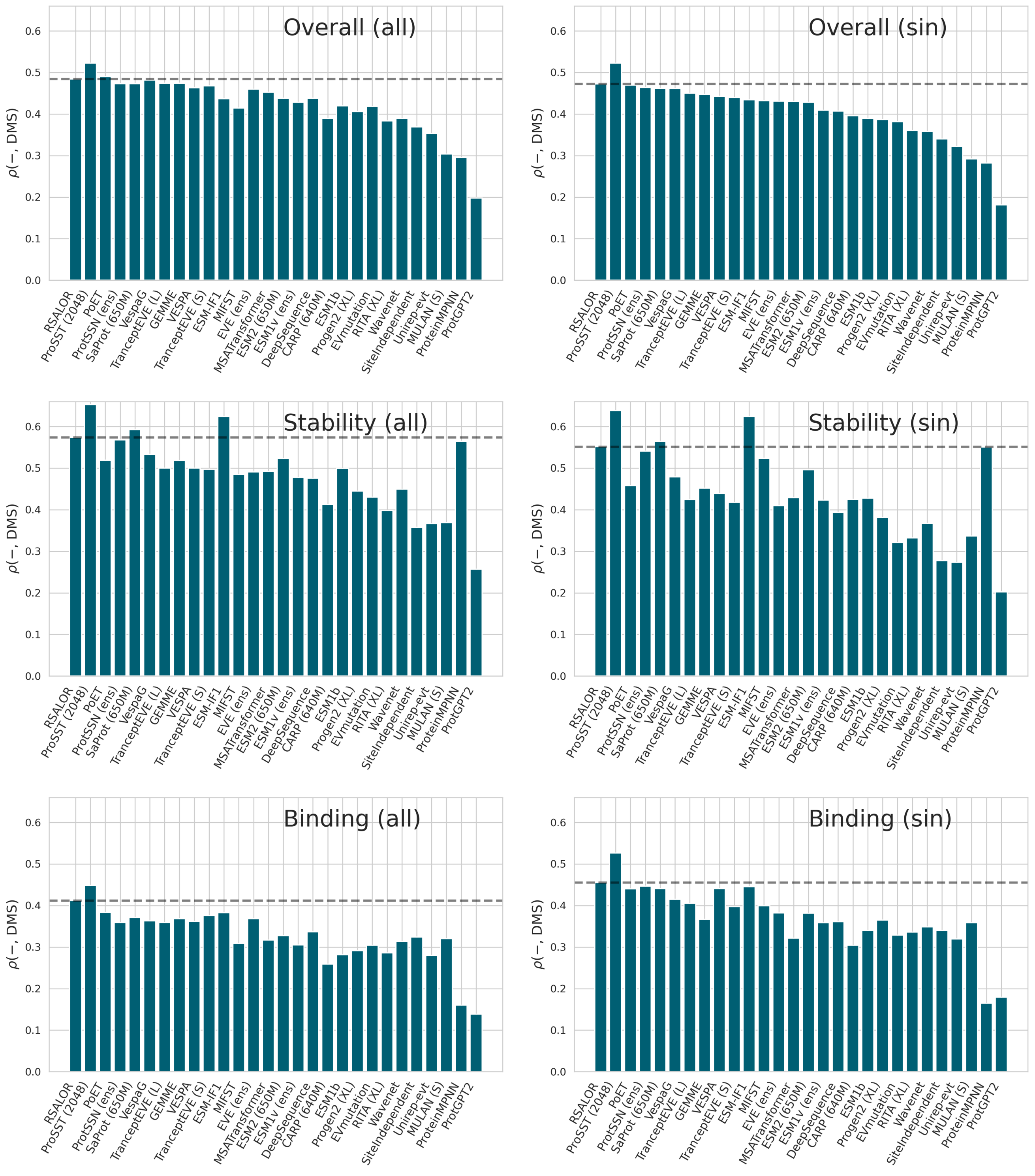

**Figure S1: Spearman correlations between predicted scores and experimental DMS scores for all datasets (overall) and categorized by target properties.** Average per-DMS Spearman correlations on all mutations (on the left) and on single-site mutations (on the right) across ProteinGym overall, stability and binding DMS datasets for RSALOR and the 27 benchmarked models. The grey horizontal line represents the RSALOR performance baseline.

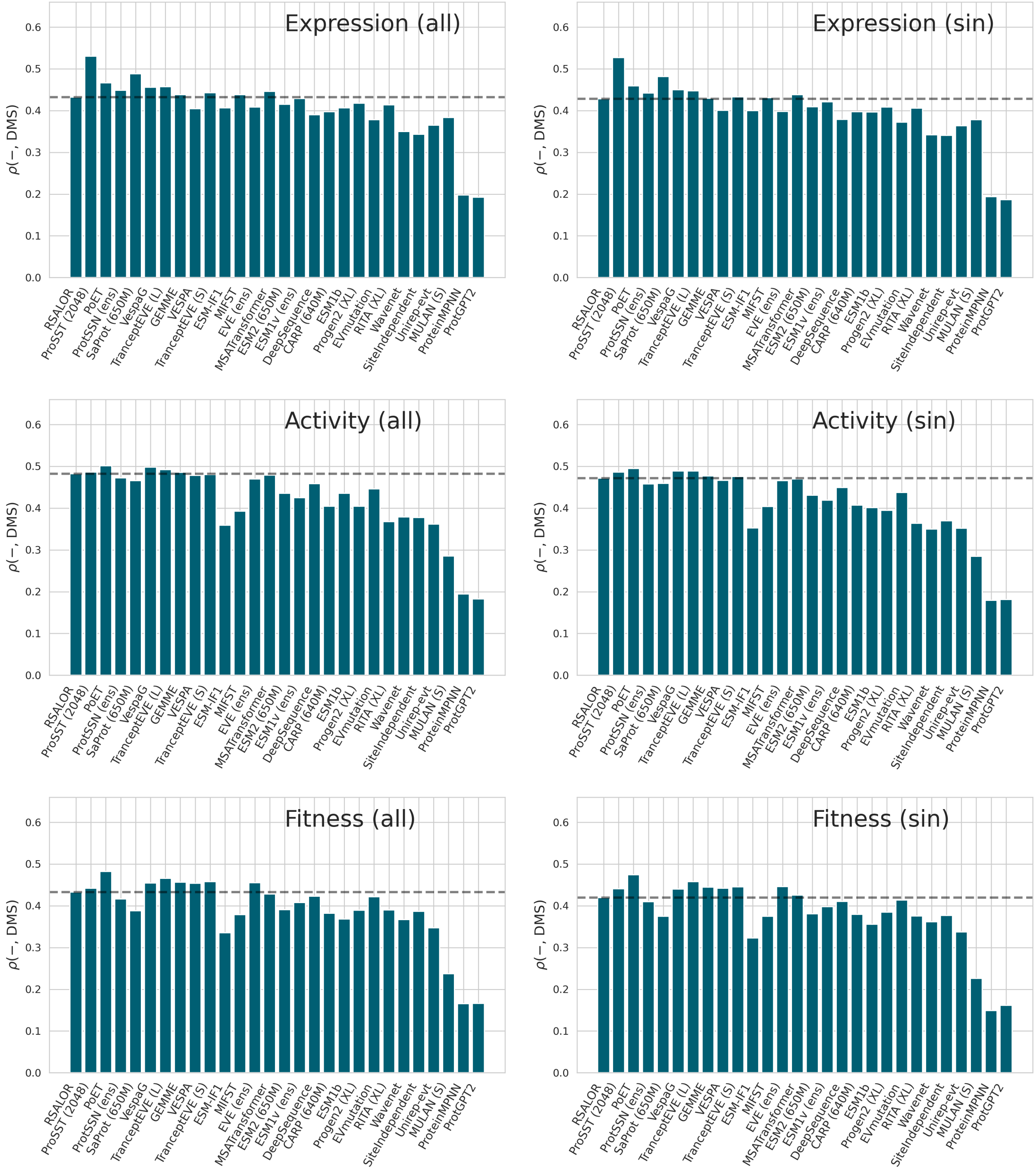

**Figure S2: Spearman correlations between predicted scores and experimental DMS scores categorized by target properties.** Average per-DMS Spearman correlations on all mutations (on the left) and on single-site mutations (on the right) across ProteinGym expression, activity and fitness DMS datasets for RSALOR and the 27 benchmarked models. The grey horizontal line represents the RSALOR performance baseline.

#### 3.2 Impact of sequence weighting

The sequence weighting process, detailed in Section 2.3, is widely used in DCA methods to correct for sampling biases caused by the selection of sequenced species (Weigt et al., 2009; Morcos et al., 2011; Ekeberg et al., 2013). In this section, we measured the impact of the weighting process on the performance of LOR and RSALOR. We showed in Tab. S4 and S5 that weighting MSA sequences mildly but consistently improves the performance of LOR and RSALOR across all DMS types and for both single-site and all mutations.

**Table S4: Impact of sequence weighting on single-site mutations**. Average per-DMS Spearman correlations across ProteinGym subclasses, comparing LOR and RSALOR models with and without using weights.

| Method | Overall | Stability | Binding | Expression | Activity | Fitness |
|---|---|---|---|---|---|---|
| LOR | 0.427 | 0.447 | 0.375 | 0.390 | 0.452 | 0.414 |
| LOR (unweighted) | 0.416 | 0.431 | 0.351 | 0.374 | 0.438 | 0.410 |
| RSALOR | 0.473 | 0.551 | 0.455 | 0.428 | 0.472 | 0.419 |
| RSALOR (unweighted) | 0.464 | 0.545 | 0.439 | 0.421 | 0.461 | 0.409 |

**Table S5: Impact of sequence weighting on all mutations**. Average per-DMS Spearman correlations across ProteinGym subclasses, comparing LOR and RSALOR models with and without using weights.

| Method | Overall | Stability | Binding | Expression | Activity | Fitness |
|---|---|---|---|---|---|---|
| LOR | 0.452 | 0.508 | 0.372 | 0.395 | 0.463 | 0.426 |
| LOR (unweighted) | 0.441 | 0.494 | 0.344 | 0.379 | 0.448 | 0.422 |
| RSALOR | 0.484 | 0.574 | 0.412 | 0.432 | 0.483 | 0.433 |
| RSALOR (unweighted) | 0.474 | 0.567 | 0.392 | 0.425 | 0.470 | 0.422 |

#### 3.3 Dependency on MSA depth

Evolutionary, sequence-based predictors tend to be more accurate when the MSA contains more sequences (Hermans et al., 2024; Laine et al., 2019; Hopf et al., 2017; Ovchinnikov et al., 2017). Indeed, a large number of homologous proteins provides a more reliable description of the mutational landscape by reducing noise in the inferred model. As in our previous work (Hermans et al., 2024), we can measure the strength of this relationship by comparing the prediction performance on each DMS dataset with the number of effective sequences in the corresponding MSA ($N_{\text{eff}}$).

Surprisingly, as shown in Fig. S3, RSALOR displays only a very weak dependency on the MSA depth, with a Spearman correlation of 0.12 between $N_{\text{eff}}$ and the model's performance on single-site mutations. Notably, the RSALOR model still provides reliable predictions even for proteins with relatively small MSAs. Very similar results were obtained on all (single-site and multiple) mutations.

In conclusion, while the relationship between MSA depth and the performance of evolutionary predictors is generally found to be stronger, the large variability in the collected DMS experiments of ProteinGym (such as DMS target property, quality and type of assay, protein size, sparsity of measured mutations) appears to have a dominant impact on prediction performance compared to $N_{\text{eff}}$.

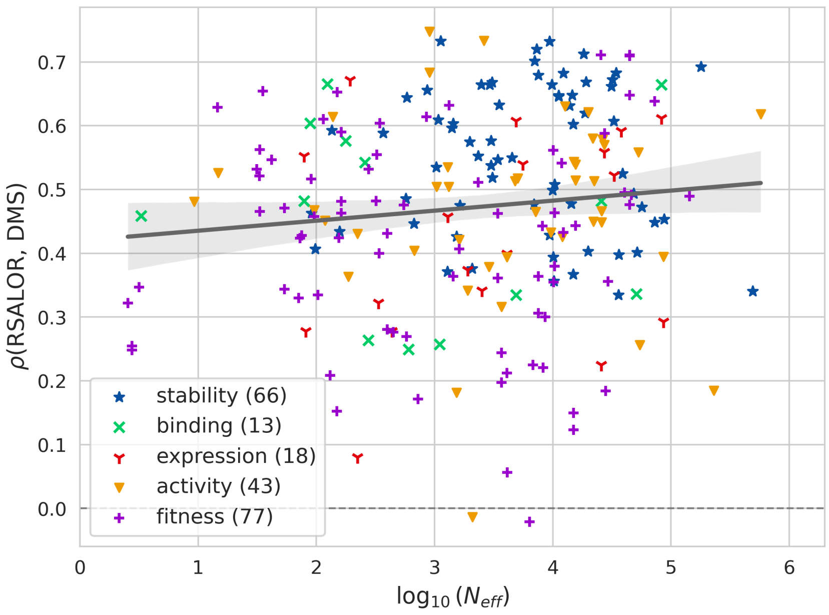

**Figure S3: Relation between performance and $N_{\mathbf{eff}}$.** Per-DMS Spearman correlations of RSALOR on single-site mutations from ProteinGym as a function of the $\log_{10}(N_{\text{eff}})$ of the corresponding MSA. Color and marker shape specify the DMS target property and the grey line represents the linear regression line.

#### 3.4 Symmetric properties of RSALOR

Structure-based predictors of the impact of mutations on protein stability and binding affinity often lack symmetry under reverse mutations (Pucci et al., 2018; Usmanova et al., 2018; Tsishyn et al., 2024b). Namely, they tend to fail to correctly predict that the effect of a mutation from states $wt$ to $mt$ is opposite to the effect of the mutation from $mt$ to $wt$. This is partly caused by the fact than they usually take as input only the structure considered as the "wild-type" while the "mutant" structure need to be somehow represented in the model. The RSALOR model also uses only the wild-type structure and computes the RSA of the mutated residue in its wild-type form. This could potentially break symmetry; therefore, we tested the model's symmetric properties here.

For this purpose, we selected one of the proteins from the ProteinGym dataset: the cellular tumor antigen p53. This protein was chosen due to its well-known role as a tumor suppressor in humans and because ProteinGym includes four DMS experiments conducted on this protein. For all possible single-site mutations from $wt$ to $mt$ on p53, we first compute the RSALOR. We then compute the RSALOR of the inverse mutations, from $mt$ to $wt$, referred to as RSALOR$^{mt}$. For that purpose, we compute the RSA of each mutated residue using the corresponding mutant structure modeled by FoldX (Delgado et al., 2019) (taking the initial wild-type structure as the template). The evolutionary component, LOR, on the other hand, is perfectly symmetric with respect to the exchange of $mt$ and $wt$.

We then examine the relationship between RSALOR and RSALOR$^{mt}$, as shown in Fig. S4. The Spearman correlation between them is $-0.976$, whereas a perfectly symmetric model would yield a correlation of $-1.00$. Moreover, regarding the impact on the performance to predict the effect of mutations, RSALOR$^{mt}$ and RSALOR display very similar correlations on the four p53 datasets (see Tab. S6). This shows that the RSALOR model is highly robust with respect to symmetry, and that using the wild-type structure as the sole structural input does not significantly impact either symmetric properties or overall performance.

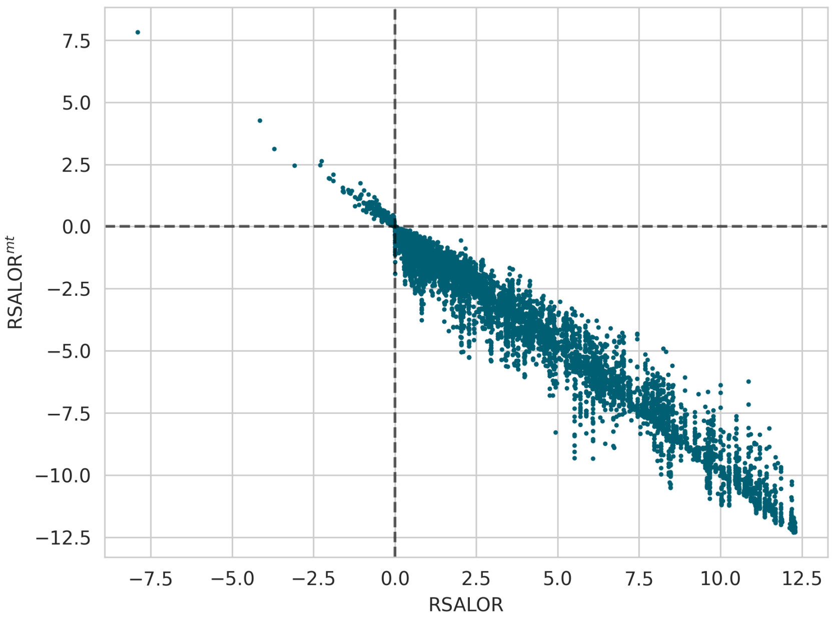

**Figure S4: Symmetric properties of RSALOR.** Comparison between RSALOR and RSALOR$^{mt}$ values on all the 7467 possible single-site mutations in the p53 structure taken from ProteinGym.

**Table S6: RSALOR and RSALOR$^{mt}$ performances**. Spearman correlations of LOR, RSALOR and RSALOR$^{mt}$ with the DMS scores on the four DMS datasets from ProteinGym that concern the p53 protein, as well as the average correlations. The DMS scores of the inverse mutations are assumed to be equal to the negative of the scores of the corresponding direct mutations. The four datasets are referenced by their ID given in the ProteinGym repository; they all contain only single-site mutations.

| **DMS dataset** | **LOR** | **RSALOR** | **RSALOR$^{mt}$** |
|---|---|---|---|
| P53_HUMAN_Giacomelli_2018_Null_Etoposide | 0.427 | 0.481 | 0.484 |
| P53_HUMAN_Giacomelli_2018_Null_Nutlin | 0.383 | 0.463 | 0.458 |
| P53_HUMAN_Giacomelli_2018_WT_Nutlin | 0.491 | 0.590 | 0.577 |
| P53_HUMAN_Kotler_2018 | 0.620 | 0.653 | 0.668 |
| **Average** | 0.480 | 0.547 | 0.547 |

#### 3.5 Impact of input MSA and structure

For the sake of fairness in benchmarking against other models referenced in the ProteinGym repository, we used the MSAs and the structures provided by ProteinGym (Notin et al., 2024). Since the target sequence of a DMS experiment is most of the time only partially covered by an experimental structure, the structures provided were generated by AlphaFold (Jumper et al., 2021). In this section, we showed that RSALOR's performance remains mostly unchanged when using different input MSAs or structures.

In our previous work (Hermans et al., 2024), we proposed a scheme to generate MSAs from the UniRef90 sequence database (Suzek et al., 2015) using the JackHMMER software (Finn et al., 2011) with 2 iterations and an $e$-value of $10^{-7}$. As shown in Tab. S7, both LOR and RSALOR exhibit almost identical performance when evaluated using the new MSAs derived with this scheme. Very similar results are observed on all (single-site and multiple) mutations.

We also tested our model on structures from the AlphaFold Protein Structure Database (AFDB) (Varadi et al., 2024), retrieved using the UniProt ID of the target sequence. While these structures are generated by the same software, they often cover a different (usually larger) range of the protein sequence and sometimes display a few residues that differ from the target sequence of the DMS. As shown in Tab. S8, RSALOR display nearly identical performance using these structures. Very similar results are observed on all (single-site and multiple) mutations.

Notably, 31 of the retrieved structures have 1 to 14 mismatched residues when aligned to those provided by ProteinGym. However, the RSALOR model is still able to make the prediction, even when the structure and the target sequence of the MSA do not perfectly match, yielding average per-DMS Spearman correlations of 0.475 and 0.471 for ProteinGym and AFDB structures, respectively. We attribute this to the fact that our model is not very sensitive to small changes in the input structure and thus does not require a high-quality experimental structure to perform well.

**Table S7: Performance using JackHMMER MSA**. Average per-DMS Spearman correlations across ProteinGym subclasses on single-site mutations, comparing LOR and RSALOR models using MSAs provided by ProteinGym and those generated with JackHMMER.

| **Method** | **Overall** | **Stability** | **Binding** | **Expression** | **Activity** | **Fitness** |
|---|---|---|---|---|---|---|
| LOR | 0.427 | 0.447 | 0.375 | 0.390 | 0.452 | 0.414 |
| LOR (JackHMMER) | 0.429 | 0.446 | 0.387 | 0.404 | 0.448 | 0.416 |
| RSALOR | 0.473 | 0.551 | 0.455 | 0.428 | 0.472 | 0.419 |
| RSALOR (JackHMMER) | 0.477 | 0.564 | 0.453 | 0.441 | 0.471 | 0.419 |

**Table S8: Performance using AFDB, the AlphaFold Protein Structure Database**. Average per-DMS Spearman correlations across ProteinGym subclasses on single-site mutations, comparing the RSALOR model using AFDB and ProteinGym structures. This table includes only the 176 DMS datasets for which an AFDB structure exists and covers all mutated residues of the DMS.

| **Method** | **Overall** | **Stability** | **Binding** | **Expression** | **Activity** | **Fitness** |
|---|---|---|---|---|---|---|
| RSALOR | 0.472 | 0.547 | 0.450 | 0.433 | 0.465 | 0.414 |
| RSALOR (AFDB) | 0.467 | 0.526 | 0.457 | 0.436 | 0.465 | 0.419 |

#### 3.6 Impact of using alternative conformations

The choice of the input conformation from which RSA is derived is important and may depend on the target biophysical property. While in our benchmark we rely on the ProteinGym-curated dataset, which consists of AlphaFold-modeled monomeric structures, predicting, for example, the effect of mutations on the binding affinity between two proteins would be more appropriate using the structure of their complex. Here, we show that one can indeed improve RSALOR performance using a more appropriate 3D conformation of the protein-protein interaction (PPI).

More specifically, we consider the DMS dataset `SPIKE_SARS2_Starr_2020_binding`, which measures the effect of mutations in the SARS-CoV-2 spike protein on its binding to human ACE2. We compare the performance of RSALOR using the AlphaFold-predicted structure of the spike protein versus the structure of the bound conformation (the heterodimeric complex available as PDB ID `6m0j` (Wang et al., 2020)). Using the experimental bound conformation substantially improves the Spearman correlation of RSALOR from 0.484 to 0.573 when restricted to the 97% of mutations located in structurally covered regions.

A similar result is observed on the dataset `RASK_HUMAN_Weng_2022_binding-DARPin_K55`, which measures the effect of mutations in the human GTPase KRas on its binding to DARPin K55 (PDB ID `5mla` (Guillard et al., 2017)). Using the experimental bound conformation, which covers 91% of the mutations, increases the Spearman correlation from 0.523 to 0.577, when restricted to structurally covered regions.

In conclusion, the choice of a more appropriate input conformation improves prediction accuracy. Mutations located at the protein–protein interface exhibit relatively high RSA values in the monomeric conformation, leading to smaller RSALOR mutational scores despite their critical role in binding affinity.

In contrast, these residues have reduced RSA in the bound conformation, resulting in higher RSALOR scores that better align with the DMS measurements.

We expect similar improvements when using 3D conformations that more closely reflect the specific biophysical property. For example, incorporating active/inactive conformations may be important when studying enzymatic properties, or using ligand-bound conformations when evaluating the impact of mutations on small-molecule binding.

### 4 Combine RSA with other models

In our previous work (Hermans et al., 2024), we showed that the use of RSA can significantly improve the performances of alignment-based models in predicting changes in protein thermodynamic stability. In this section, we show that this statement can be generalized to a much wider range of models and to various types of mutational effects. The results are presented on LOR (which defines our RSALOR model) and on the 27 benchmarked models (defined in Section 3.1).

Impressively, as shown in Fig. S5, combining predictions with RSA, using Eq. 10, substantially improves the performance of nearly all models. This holds true even for models that explicitly use the structure as input (labeled in Tab. S1).

We believe that, in the future, incorporating RSA and other types of structural knowledge for mutational landscape predictions could be a valuable source of information for a wide range of model types. Although the use of RSA to predict the effect of mutations is not new, this observation further highlights the importance of integrating simple structural information with other types of data to enhance models' performance.

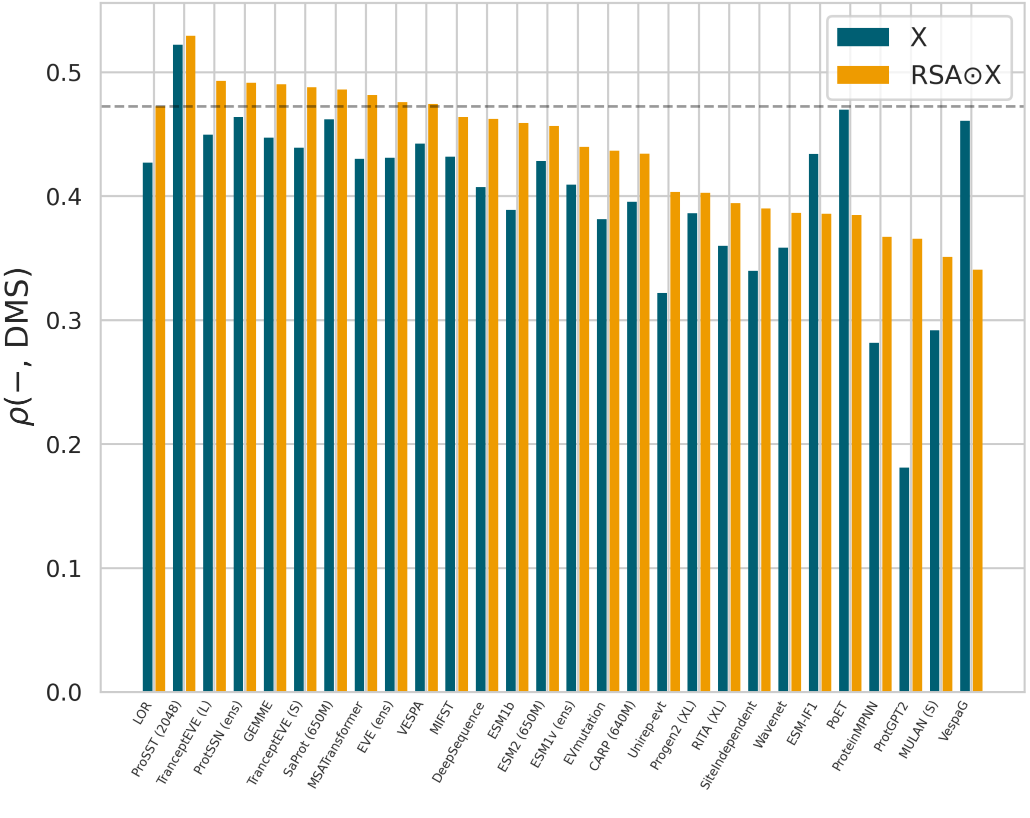

**Figure S5: Using RSA with other models.** Average per-DMS Spearman correlations on single-site mutations from ProteinGym for LOR and the 27 benchmarked models (in blue), as well as their combination with RSA (in orange). The grey horizontal line represents the RSALOR performance as a baseline.
